# spatialgater: A Shiny webtool for *in situ* gating of cells in SpatialExperiments

**DOI:** 10.1101/2025.10.26.684544

**Authors:** Markus Steiner, Stephan Drothler, Jan Philip Höpner, Roland Geisberger, Nadja Zaborsky

## Abstract

Multiplexed imaging techniques generate high-dimensional datasets that contain their molecular profiles of cells combined with spatial coordinates, which can be stored in `SpatialExperiment` objects.

Current analysis workflows separate cells after clustering them by their bio-molecule expression levels without considering their spatial context within the tissue. While patch-/neighbourhood detection methods exist, there is no option to select single cells by their location at will. By introducing `spatialgater`, we aim to boost interactivity and reduce programming efforts of image analysis by enabling spatial gating of cells from `SpatialExperiment` objects through an intuitive web-based interface. The package displays cells as dots on a zoom-able image and allows users to draw polygon gates directly on individual cells. All selected cell identifiers can be exported as.csv and/or written back as logical gate column in the `colData()` of the original `SpatialExperiment`.

`spatialgater` provides an accessible, interactive addition to spatial analysis pipelines. By using a publicly available imaging mass cytometry dataset from breast cancer tissue, we demonstrate its effectiveness in characterizing and comparing T-cells by their spatial location.

## I. INTRODUCTION

Spatial “omics” technologies have rapidly expanded the biological questions that can be addressed *in situ*, particularly in cancer during recent years. Hereby, analysis of specific cell phenotypes, cell states and precise location of single cells in tissue help in understanding of complex diseases on a molecular level [1–4].

Within the R/Bioconductor ecosystem, the S4-vector based `SpatialExperiment` extends the `SingleCellExperiment` class with coordinates and image data for downstream analysis. It is one of the most versatile data container formats used for imaging mass cytometry data. It harbors many important visualization and analysis functions and packages which extend its range of application even further [5–7].

The `cytomapper` package focuses on visualization of pixel-level and cell-level information on segmentation masks and supports mapping of interactively gated cells from expression dot-plots back to the images via `cytomapperShiny()` [8]. Similarly, `cytoviewer` provides an interactive visualization tool for channel data, segmentation masks, and `colData()` entries [9]. Thus, both tools provide powerful and complementary visualization options, but lack the ability to select certain cell populations by location or spatial context.

Here, we present the R package: `spatialgater`, an R/Shiny tool that can add to the analysis workflow by enabling cell-selection by location (theoretically down to a single cell). The user is able to draw gates on cell coordinates to capture anatomically or histologically striking regions, and the gate assignment is transferred as logical column to the `SpatialExperiment` for immediate use in downstream analysis. This approach integrates well with the Bioconductor workflow and requires minimal programming, in contrast to traditional ways of spatial community or patch detection analysis [7]. In sum, interactive spatial gating adds a lightweight yet effective layer to spatial omics pipelines, improving interpretability and accelerating exploratory-to-confirmatory analysis. Here we show the application of `spatialgater` on an imaging mass cytometry data set of breast cancer tissue. Using `spatialgater()` we select T-cells close to patches of CK7^+^CK^+^ and CK^+^HRhi tumor cells and compare protein expression to T-cells, which are located further away from these tumors clusters.

## II. METHODS

### A. Software Architecture

The R package `spatialgater` is built on the Shiny web app framework for R [11]. The package consists of three main components: (1) data processing of `SpatialExperiment` objects, (2) the Shiny user interface for interactive visualization and gating and (3) export mechanisms that integrate the gated cell IDs back into the original data structure with optional CSV export. Installation instructions are described in the github repository (see section VI).

#### 1) Dependencies

The app depends on several key R packages:

- `SpatialExperiment`: Provides the core data structure for spatial omics data [12]
- `Shiny`: Enables interactive web application [11]
- `leaflet/leaflet.extras`: Provides interactive mapping with polygon drawing tools [13, 14]
- `sf`: Handles spatial geometry operations for point-in-polygon calculations [15]

#### 2) User Interface

The interface consists of three main components:

1. Control Panel: Contains sample selection dropdown, gate naming input, action buttons for export, download, and clearing of gated cell IDs
2. Interactive Map: Displays cell positions as colored dots and a color legend. Users can zoom and draw polygons on the map.
3. Selection Display: Shows the current selection of cell IDs as a union of all active polygons from a single sample. These IDs will be exported. Zooming option allows for gating of a single cell.

#### 3) Coordinate and Gate handling

The first two columns of `spatialCoords()` are read as pixel coordinates. The Y-axis is mirrored to replicate the image-style coordinates. Visualization and point-in-polygon queries run in this mirrored space. Drawn polygons are non-permanent: the app maintains only the union of cell IDs currently covered by the gates for the selected sample. Exporting is “add-only” and writes logical gate columns to `colData(spe)`$`<gate> <-TRUE` for those cells. It does not reset prior `TRUE` values to `FALSE`. The same cell IDs can be downloaded as a csv. Switching samples clears polygons and the live selection but preserves any previously exported gate columns. Optionally, a user-supplied `on_export()` callback can override writing. If enabled, `auto_write_back` assigns the updated `SpatialExperiment` object to the calling environment when the `spatialgater` session ends.

### B. Analysis Procedure

Rstudio (V 4.5.1) equipped with the following packages was used to conduct the analysis [16–23]:

**library** (“ s p a t i a l g a t e r “)

**library** (“ h e r e “)

**library** (“ i m c d a t a s e t s “)

**library** (“ t i d y v e r s e “)

**library** (“ s v g l i t e “)

**library** (“ RColorBrewer “)

**library** (“ ggpubr “)

**library** (“ scater “)

Statistical tests were performed using `geom_pwc()` with standard parameters (wilcox-test). UMAP was calculated using `runUMAP()`. Plots were generated using `ggplot2` and `dittoSeq` and assembled with the help of `magick`, `grid` and `patchwork` [19, 24–27]. Statistical significance was assumed at p*<*0.05. T-cell groups were defined by visual approximation. Tumoral T-cells were gated if not more than roughly 4 cells away from a CK7^+^CK^+^ and CK^+^HRhi tumor patches. Distal T-cells were gated if they appeared at least 10 cells away from a CK7^+^CK^+^ and CK^+^HRhi tumor patch. Tumor cells were assumed as patches if there were at least 5 tumor cells in direct cell-cell contact. Among all T-cells in the sample (image id: 156), we gated an exemplary region of 135 tumoral T-cells and 99 distal T-cells. A link to the analysis script can be found in section VI.

## III. RESULTS

### Convenient characterization of tumor-adjacent T-cells

To test the functionality of `spatialgater` we downloaded a public imaging mass cytometry data set of breast cancer tissue by Jackson et al. sourced via the `imcdatasets` package [10, 18]. The data set (with standard import parameters) comprises 100 samples of breast cancer tissue, analyzed with 35 isotope-conjugated antibodies. The authors identified 27 metaclusters falling into the following four categories: immune, epithelial, stromal and tumor. We were interested if there are phenotypical differences between T-cells which are located close to tumor cells (approx. not more than four cells away from a CK7^+^CK^+^ and CK^+^HRhi tumor patch) which we defined as tumoral T-cells, compared to “distal” T-cells which are located further away (approx. more than ten cells away from a CK7^+^CK^+^ and CK^+^HRhi tumor patch).

To begin with, we screened the `SpatialExperiment` for samples with high amounts of T-cells and selected a sample (image id: 156) based on visual inspection with following selection parameters:

1. Presence of distinct tumor cell patches
2. Presence of T-cells close to these tumor patches
3. Presence of T-cells distant from these tumor patches

Next, we called `spatialgater()` and gated 135 tumoral T-cells and 99 distal T-cells (see. Fig 1). We then proceeded to investigate the phenotype of these T-cells.

**Figure 1.**
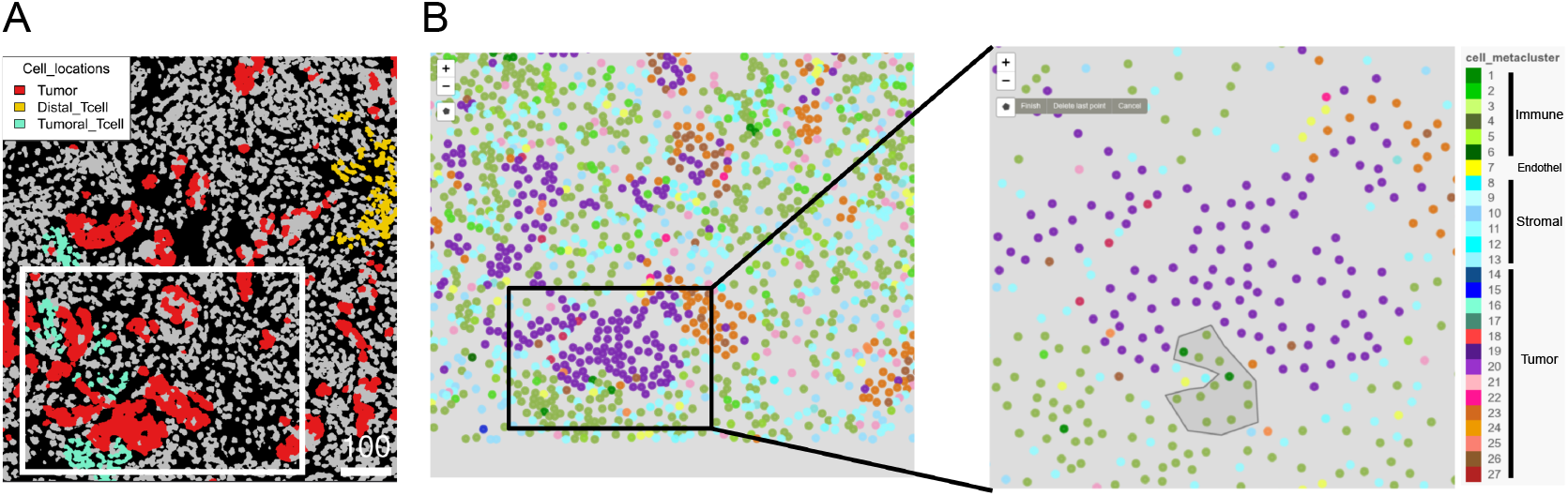
Gating of T-cells close and distant to breast cancer cells from the Jackson et al. data [10]. **A**: Final gating strategy visualized using cell level plots of the original cell masks using `cytoviewer()` [9]. Tumor cells are highlighted in red and are derived from cell metacluster 20 and 23 (see legend in Figure 1 B) which are defined as CK7^+^CK^+^ and CK^+^HRhi by Jackson and colleagues. T-cells close to tumor patches are highlighted in aquamarine (`Tumoral_Tcell`), while T-cells further away from tumor patches are highlighted in gold (`Distal_Tcell`) (definition of cell types in section II-B). All other cell masks are colored in grey. Scale bar (right bottom) is shown in µm. The white square marks the gating location highlighted in Figure 1B. **B**: Crop of the `spatialgater` user-interface (image id: 156) of the data set (left image), approximately marked by the white rectangle in Figure 1A. Zoom-in image, demonstrating polygon-based gating on a small population of tumoral T-cells (grey shaded, cell metacluster: 3, 5) in close proximity to CK7^+^CK^+^ tumor cells (cell metacluster: 20) (right image).

First we calculated a Uniform Manifold Approximation and Projection (UMAP) of all cells in image 156 and localized distant and tumoral T-cells (see. Figure 2A-C). Tumoral and distal T-cells seem to cluster slightly different within the immune cell compartment (green clusters, cell metacluster: 1-6), indicating differences in expression profiles. Importantly, tumoral T-cells did not cluster near tumor cells in the UMAP, demonstrating that T-cells did not suffer from lateral spillover of neighboring cells. Finally we compared expression levels of the state markers CD44, GATA3, H3, Ki67 and phosphorylated mTOR (p mTOR) between the two populations. T-cells close to the tumor patches had a significantly higher expression of activation markers CD44, GATA3, H3 and p mTOR (see. Figure 2D, p*<*0.001), which indicates ongoing immune activation and potential localized anti-tumor immune responses.

**Figure 2.**
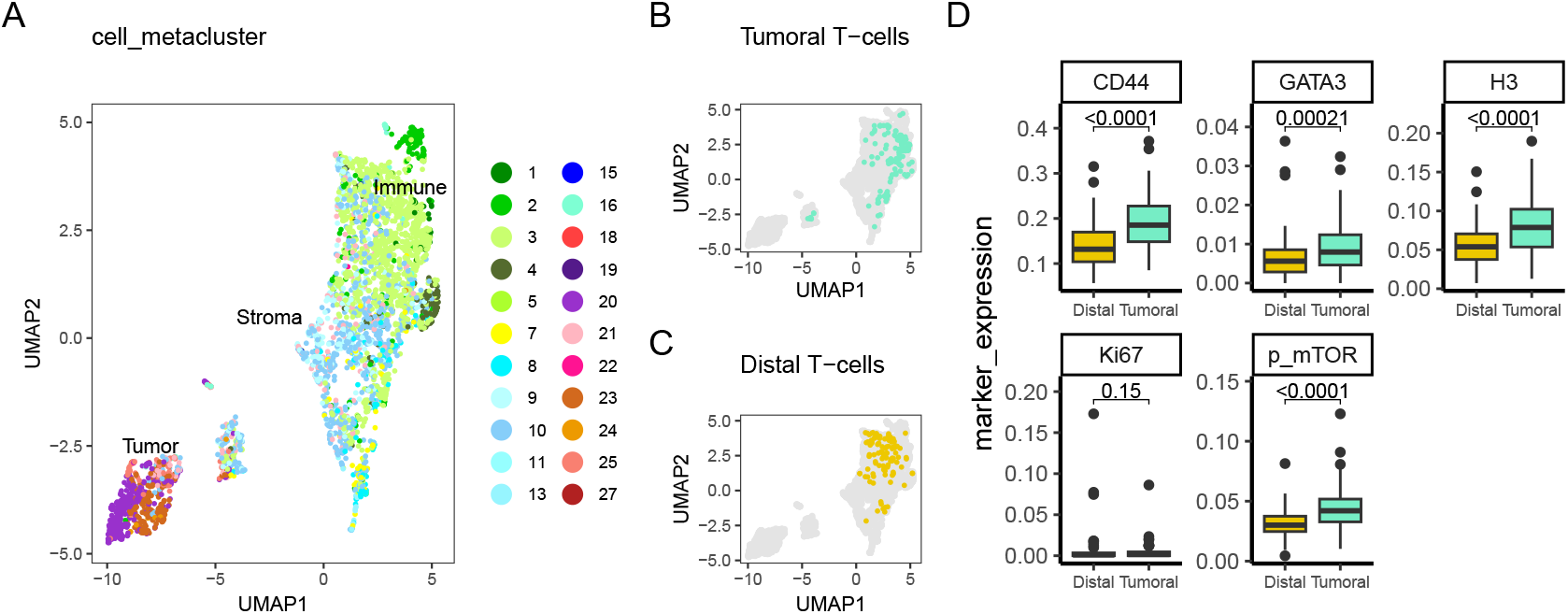
Distinct clustering and marker expression of tumoral and distal T-cells from beast cancer sample (image id: 156) [10] **A**: UMAP of all cells from the sample (image id: 156) with annotations of “Immune” (cluster: 1-5), “Stroma” (cluster: 8-13) and “Tumor” cells (cluster: 15-27). **B**: UMAP, highlighting tumoral T-cells (T-cells, close to patches of tumor cells, see. 1B), while remaining cells are colored grey. **C**: UMAP, highlighting distal T-cells (T-cells, located away from tumor patches, see. 1B), while remaining cells are colored grey. **D**: Boxplots of state marker expression (@assay = quant_norm) of distal and tumoral T-cells. Statistical comparison conducted via wilcoxon test, significance assumed at p*<*0.05. *Abbrev:* p mTOR: phosphorylated mTOR

## IV. DISCUSSION

We introduced `spatialgater`, a Shiny webtool, which allows for fast and convenient gating of cells using spatial information. Cells are displayed as dots, positioned by their coordinates from the imaging data while gating is realized by a polygon-based selection strategy. Thereby, the program is lightweight and easy to navigate, while providing a useful add-on to existing image analysis pipelines.

We demonstrated the functionality of the app by downloading a public data set of breast cancer tissue, measured by imaging mass cytometry. We used `spatialgater` to select two populations of T-cells in one of the samples (image id: 156) from the data set. For the first population, T-cells were gated if they were in close proximity to patches of tumor cells (tumoral T-cells), while the second group of T-cells was selected based on a larger distance to tumor patches (distal T-cells) (described in section II-B).

We compared protein expression of several state markers between the two groups of T-cells and found higher activation in T-cells which are in close contact to tumor cells. This complements the current state-of the art in image analysis, where dimensionality reduction, clustering and patch detection algorithms are applied to the cell data. These clustering outcomes often depend on user input, are frequently resource intensive and outcome optimization requires time consuming trial-and-error approaches. `spatialgater` provides an alternative way of selecting cells, which might be of interest solely based on their location within tissues, without the need of discriminating them by computational approaches.

In its current form, `spatialgater` is optimized and tested on imaging mass cytometry data, where regions of interest (ROI) are usually quite small compared to immune fluorescence and immune histochemistry. Since crowding effects (e.g. overlay of dots at low zoom levels) and performance limitations (e.g. large ROIs filling up random-access memory) employed by other imaging techniques, cannot be ruled out, `spatialgater` might be updated in regards to zooming, ROI navigation and processing capabilities in the future.

Since the position of a cell in a tissue is a prerequisite to perform its intended function, `spatialgater` can be useful not only in tumor immunology, but also in other branches of life sciences.

In summary, we provide a compact and useful Shiny tool, which can be easily implemented in already established analysis workflows of imaging data, by providing simple and swift selection of specific cells or cell types based on their location *in situ*.

## VI. DATA AVAILABILITY

The github repository for `spatialgater` and scripts used to produce the findings in this paper are available under: https://github.com/Mark-Ste/spatialgater

## VII. AUTHOR CONTRIBUTIONS

MS wrote the app, reviewed data analysis and wrote the manuscript. SD analyzed data, reviewed the app and wrote the manuscript. JPH curated the data and supported app development. RG conceptualized the project, supported app development and revised the manuscript. NZ conceptualized and supervised the project, supported app development, reviewed data analysis and revised the manuscript. All authors read, reviewed and agreed to the final version of the manuscript.

## VIII. ACKNOWLEDGMENT

The authors thank Anna Kolmhofer, Christian Scherhäufl for their help in figure conceptualization and proofreading.

